# Deuteros: software for rapid analysis and visualization of data from differential hydrogen deuterium exchange-mass spectrometry

**DOI:** 10.1101/417998

**Authors:** Andy M. C. Lau, Zainab Ahdash, Chloe Martens, Argyris Politis

**Author notes:** These authors contributed equally (Zainab Ahdash, Chloe Martens).

## Abstract

**Summary:** Hydrogen deuterium exchange-mass spectrometry (HDX-MS) has emerged as a powerful technique for interrogating the conformational dynamics of proteins and their complexes. Currently, analysis of HDX-MS data remains a laborious procedure, mainly due to the lack of streamlined software to process the large datasets. We present Deuteros which is a standalone software designed to be coupled with Waters DynamX HDX data analysis software, allowing the rapid analysis and visualization of data from differential HDX-MS.

**Availability:** Deuteros is open-source and can be downloaded from https://github.com/andymlau/Deuteros, under the Apache 2.0 license.

**Implementation:** written in MATLAB and supported on both Windows and MacOS. Requires the MATLAB runtime library.

## Introduction

Hydrogen deuterium exchange mass spectrometry (HDX-MS) is a structural technique which has garnered attention for its ability to assess protein-protein and protein-ligand interactions, protein folding, and the associated dynamics of these processes (Jensen, 2016; Konermann, et al., 2011; Masson, et al., 2017; Mistarz, et al., 2016). The basis of HDX-MS relies on the exchange of labile amide hydrogens of the protein backbones, for bulk deuterium within solution. The protein of interest is allowed to undergo exchange in a deuterium-rich buffer for a set number of timepoints and then quenched. The protein is then enzymatically cleaved to the peptide level, and the mixture is subjected to liquid chromatography coupled to mass spectrometry (LC-MS). Using LC-MS, the mass of the peptide acquired through deuteration can be determined via a database search. Peptides which participate in hydrogen bonding of amide hydrogens result in lesser exchange (Marcsisin and Engen, 2010). Additionally, those which comprise the accessible surfaces of proteins, may experience relatively greater deuteration, than those found in the protein interior (Marcsisin and Engen, 2010). In differential HDX-MS (ΔHDX-MS), peptides from a reference state are compared with those from an altered state (which could be for instance a mutation or a ligand) to report on regions of the protein which are affected by structural or conformational perturbations.

While automated data processing software exist for data analysis of raw HDX-MS data (Claesen and Burzykowski, 2017; Guttman, et al., 2013; Pascal, et al., 2012; Rey, et al., 2014), visualization of the results prior to interpretation can be challenging due to the size of the datasets involved. A typical HDX-MS experiment will result in the order of 10^2^ peptides depending on the system size and complexity. Several visualization methods have been introduced to provide clarity on HDXMS data interpretation. These can be divided into graphical representations including the Woods plot (Woods and Hamuro, 2001) and uptake maps, and molecular representations where uptake and other data are projected onto 3-dimensional structures of the system (Kavan and Man, 2011).

To simplify the HDX-MS analysis workflow, we have developed Deuteros for the rapid visualization of differential HDX-MS data. Development of the software was primarily motivated by the lack of a streamlined workflow for differential data analysis and visualisation, particularly for Waters HDX-MS instrumentation. Deuteros provides a user-friendly interface for ΔHDX-MS data visualisation, which can be unnecessarily laborious for large datasets. It also provides simple statistical evaluation of peptide deuteration which can be used to identify biologically interestingly regions of proteins. Finally, inputs to Deuteros have been standardized to csv files, allowing the software to be potentially compatible with any instrumentation.

We have applied our software to a comparison of the wild-type xylose transporter (XylE) and a E153Q mutant. XylE is a secondary membrane transporter protein tasked with the role of shuttling xylose sugar across bacterial cell membranes (Quistgaard, et al., 2013). A member of the Major Facilitator Superfamily (MFS), XylE operates through an alternating-access mechanism, transitioning between inward-facing and outward-facing conformational states in a highly dynamic fashion (Wisedchaisri, et al., 2014).

## How does it work?

Deuteros is a standalone MATLAB application available to both MacOS and Windows and requires the MATLAB runtime library. The application requires two inputs: the ‘state’ and ‘difference’ files exported from DynamX HDX data analysis software (Waters Corp.). Deuteros analysis consists of four steps including data input and three visualisation stages: flattened data maps, Woods plot (with statistical peptide filtering) and output to PyMOL.

### Input data

The DynamX ‘state’ file contains a per-protein, per-peptide, per-time point aggregation of peptide deuterium uptake data from the ΔHDX-MS conditions. State files contain information including m/z, maximum possible deuterium uptake, observed deuterium uptake, standard deviation, retention time, and any residue modifications reported. Users should only enable proteins and states of interest and disable all others within the DynamX session file. The ‘difference’ file contains a per-peptide, per-timepoint comparison of peptide deuterium uptake from two user defined states. The difference file can only be generated from DynamX when two or more states are loaded into the dataset. Users should ensure that the correct comparison is made by selecting the correct states within DynamX. A video tutorial and example datasets have been provided alongside the software.

### Visualization

Deuteros produces flattened data maps including coverage, residue-level redundancy, deuterium uptake heat maps and Woods plots. Coverage, redundancy and various deuterium uptake styles can also be exported from Deuteros, to be projected onto atomic models of the protein of interest in the PyMOL molecular graphics viewer (Schrödinger, 2015). Users can simply ‘drag and drop’ these files into PyMOL (for MacOS, or PyMOL version 2.0 and above for Windows), or alternatively copy and paste the contents of the file into the PyMOL command line.

### Statistics

Confidence limits are calculated as (Houde, et al., 2011):

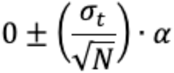

Where *σ*_*t*_ is the standard deviation of the mean uptake for timepoint *t, N* is the number of sample replicates and *α* is the critical value desired. By default, Deuteros provides critical values for 95 and 99% confidence limits (4.303 and 9.925) for a two tailed t-test with *df*=2 degrees of freedom. For ‘sum’ data, where peptide deuterium uptake differences from each timepoint are aggregated together to better identify potential peptides that are conformationally active, *σ*_*t*_ of each peptide is summed over all timepoints, while considering error propagation from each time component.

### Application

To showcase the capabilities of Deuteros, we imported state and difference files from the wild-type and E153Q mutant XylE membrane transporter. The coverage map of XylE indicate a 92.5% coverage, with the largest non-covered region around residues 230-240 (Figure 1a). The redundancy map expands on the coverage map, displaying the same coverage, but with a white-magenta color gradient to represent peptide redundancy. Reviewing the map shows that the highest redundancy of the XylE dataset was at 12 peptide copies around residues 465 (Figure 1b). The Nterminal, residues 90, 180, 260 and 360 have only a single peptide representing these regions.

**Figure 1.**
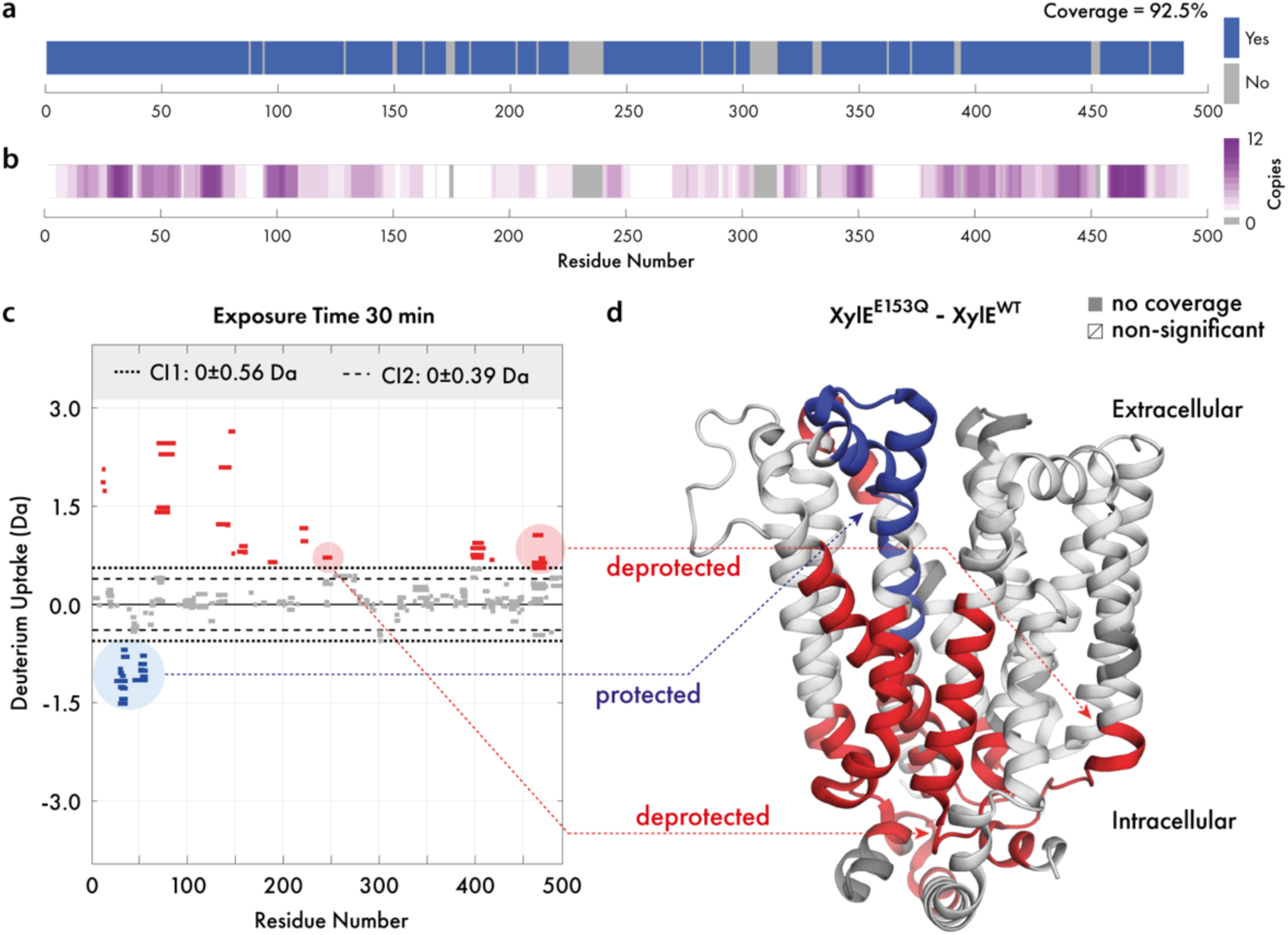
Overview of Deuteros. Visualisation of (a) experimental protein coverage, (b) data redundancy and (c) deuterium uptake differences in Woods plot format. Dashed and dotted lines indicate 95 and 99% confidence limits applied to the dataset to identify peptides with significant deuteration differences. Deprotected, protected and non-significantly different peptides are in red, blue and grey respectively. (d) Differential HDX-MS data for the wild-type and E153Q mutant XylE has been projected onto its crystal structure (PDB ID: 4GBY).

The Woods plot section displays a per-timepoint breakdown of the differential dataset in a grid layout. Deuteros can display a maximum of approximately 8 timepoints simultaneously before individual Woods plots become too crowded, depending on the screen size and resolution. Woods plots first apply confidence filtering to all peptides in each timepoint (Figure 1c). Peptides with differential deuteration outside of the user selected confidence limits are non-significant and are shown in grey. The significant peptides are shown as red for deprotected and blue for those that are protected. While only one set of confidence limits are applied to the data, two boundaries are shown on each Woods plot as a visual aid for users to view which peptides might be significant, should they wish to tighten or relax the filter used. The legend section displays the confidence limits as ± Da (to two decimal places) values around 0 (or no difference). To facilitate interpretation, significant peptides can also be exported as a *csv* file containing a per-peptide per-timepoint breakdown of the ΔHDX-MS data. Users may also take advantage of the in-built MATLAB data cursor which displays the residue number and differential uptake of a peptide by clicking on the peptide within the graphical user interface.

The PyMOL export section consists of options for formatting the data from the linear coverage map and Woods plot sections, for visualization in PyMOL through *pml* files. Coverage and redundancy can be projected onto structures and a range of color palettes are available. Differential deuteration data can also be exported for projection onto the molecular structure of XylE (PDB ID: 4GBY; Figure 1d). For this representation, the deuteration data type can show absolute differential uptake (in Daltons), or the differential relative fractional uptake (ΔRFU). The ΔRFU considers the peptide length and its maximum deuteration and scales the absolute uptake as a percentage of this value, which may be more informative for some datasets. Similar to the Woods plot, Deuteros implements red/blue/white/grey color scheme for protected, deprotected, non-significant and non-covered regions. Through projection of deuteration data onto the structure of XylE, structural effects caused by the E153Q mutation are immediately visible (Figure 1d). The extracellular-facing portion of XylE experiences protection (blue), while the intracellular portion experiences deprotection (red).

## Acknowledgements

We would like to acknowledge Jurgen Claesen (Hasselt University) for critically review of the manuscript and Eamonn Reading (King’s College London) for suggestions with software features.

## References

Claesen, J. and Burzykowski, T. Computational methods and challenges in hydrogen/deuterium exchange mass spectrometry. Mass Spectrom Rev 2017;36(5):649–667.

Guttman, M., et al. Analysis of overlapped and noisy hydrogen/deuterium exchange mass spectra. J Am Soc Mass Spectrom 2013;24(12):1906–1912.

Houde, D., Berkowitz, S.A. and Engen, J.R. The utility of hydrogen/deuterium exchange mass spectrometry in biopharmaceutical comparability studies. J Pharm Sci 2011;100(6):2071–2086.

Jensen, P.F., Rand, K. D. Hydrogen Exchange. In Hydrogen Exchange Mass Spectrometry of Proteins, D. D. Weis (Ed.). doi:10.1002/9781118703748.ch1. 2016.

Kavan, D. and Man, P. MSTools-Web based application for visualization and presentation of HXMS data. Int J Mass Spectrom 2011;302(1-3):53–58.

Konermann, L., Pan, J. and Liu, Y.H. Hydrogen exchange mass spectrometry for studying protein structure and dynamics. Chem Soc Rev 2011;40(3):1224–1234.

Marcsisin, S.R. and Engen, J.R. Hydrogen exchange mass spectrometry: what is it and what can it tell us? Analytical and bioanalytical chemistry 2010;397(3):967–972.

Masson, G.R., Jenkins, M.L. and Burke, J.E. An overview of hydrogen deuterium exchange mass spectrometry (HDX-MS) in drug discovery. Expert Opin Drug Discov 2017;12(10):981–994.

Mistarz, U.H., et al. Probing the Binding Interfaces of Protein Complexes Using Gas-Phase H/D Exchange Mass Spectrometry. Structure 2016;24(2):310–318.

Pascal, B.D., et al. HDX workbench: software for the analysis of H/D exchange MS data. J Am Soc Mass Spectrom 2012;23(9):1512–1521.

Quistgaard, E.M., et al. Structural basis for substrate transport in the GLUT-homology family of monosaccharide transporters. Nat Struct Mol Biol 2013;20(6):766–768.

Rey, M., et al. Mass spec studio for integrative structural biology. Structure 2014;22(10):1538–1548.

Schrödinger, L. The PyMOL Molecular Graphics System, Version 1.8. 2015.

Wisedchaisri, G., et al. Proton-coupled sugar transport in the prototypical major facilitator superfamily protein XylE. Nat Commun 2014;5:4521.

Woods, V.L., Jr. and Hamuro, Y. High resolution, high-throughput amide deuterium exchange-mass spectrometry (DXMS) determination of protein binding site structure and dynamics: utility in pharmaceutical design. J Cell Biochem Suppl 2001;Suppl 37:89–98.

